# Age-specific walking speed during locomotor adaptation leads to more generalization across contexts

**DOI:** 10.1101/2023.08.10.552802

**Authors:** Dulce M. Mariscal, Carly J. Sombric, Gelsy Torres-Oviedo

## Abstract

Previous work has shown that compared with young adults, older adults generalize their walking patterns more across environments that impose different motor demands (i.e., split-belt treadmill vs. overground). However, in this previous study, all participants walked at a speed that was more comfortable for older adults than young participants, which leads to the question of whether young adults would generalize more their walking patterns than older adults when exposed to faster speeds that are more comfortable for them. To address this question, we examined the interaction between healthy aging and walking speed on the generalization of a pattern learned on a split-belt treadmill (i.e., legs moving at different speeds) to overground. We hypothesized that walking speed during split-belt walking regulates the generalization of walking patterns in an age-specific manner. To this end, groups of young (<30 y/o) and older (65+ y/o) adults adapted their gait on a split-belt treadmill at either slower or faster walking speeds. We assessed the generalization of movements between the groups by quantifying their aftereffects during overground walking, where larger overground aftereffects represent more generalization, and zero aftereffects represent no generalization. We found an interaction between age and walking speed in the generalization of walking patterns. More specifically, older adults generalized more when adapted at slower speeds, whereas younger adults did so when adapted at faster speeds. These results suggest that comfortable walking speeds lead to more generalization of newly acquired motor patterns beyond the training contexts.

## INTRODUCTION

Generalization is the ability to carryover learned movements from trained to untrained conditions. For example, most people learn how to swim in a controlled environment like a pool but can use the learned movements to swim in a more uncontrolled environment like the ocean. While our motor system can adapt and learn new movements with training devices, like exoskeletons or treadmills, it has been shown that not everything learned in the training environment carries over to movements without the device[1]–[4]. Understanding the factors limiting the generalization of motor learning is critical to improving the effectiveness of rehabilitation; more specifically, the corrective movements that patients learn in the clinic must generalize to their day-to-day activities. Thus, we are interested in identifying factors mediating the generalization of motor learning to harness them such that motor improvements acquired in the clinic generalize to untrained circumstances.

Previous findings indicate that context cues regulate generalization. For example, unusual movements during adaptation [4], [5] or pre-movements to the task [6]–[8] can modulate the generalization of movements. Furthermore, it has been shown that the generalization of motor memories is reduced as a function of the difference between the testing and the training conditions [9], [10] and that differences between the walking patterns on the treadmill and overground limit the generalization of movements between the two contexts [4], [11]. These findings suggest that similarities between context cues specific to the training and testing conditions would lead to greater generalization of movements beyond the training context.

In the case of locomotion, walking speed might be a contextual cue modulating the similarity between training and testing contexts. In other words, walking at unusual speeds during training (e.g., too fast or too slow) may reduce the generalization of learned movements. Previous work observed that older adults generalize more than younger individuals [12] when adapted at slow walking speeds ([0.5 – 1 m/s]) that were more comfortable for older than younger people [13]–[15], raising the possibility for walking speed to be a modulator on the distinct age-specific generalization of walking movements. Alternatively, the aging effect on generalization might be explained by an age-related decay in neural processes needed for motor switching [12], [16], irrespective of the walking speed at which participants are adapted.

In this study, we investigated the interaction between walking speed and healthy aging on the generalization of locomotor patterns (split-belt treadmill walking to overground walking). We hypothesized that walking speed modulates generalization in an age-specific manner. In other words, we expected that younger adults would generalize more when adapted to faster walking speeds, whereas older adults would generalize more when adapted to slower speeds. To test this, we adapted older and younger adults to a split-belt perturbation at either slow or fast speeds. We measured how much the adapted walking patterns generalized from the treadmill (training context) to overground (testing context). If our hypothesis is supported, our results suggest that training at self-selected walking speed leads to more generalization of movements beyond the training context.

## METHODS

We studied how healthy aging and walking speed affect the adaptation and generalization of split-belt walking patterns. To test the effect of healthy aging, 16 young adults (7 females, 20.81 ±6.02 yrs.) and 16 older adults (9 females, 73.13±5.40 yrs.) participated in a split-belt adaptation paradigm. Additionally, to test the effect of walking speed, participants from both age groups experienced the same split-belt paradigm at either slow or fast mean walking speeds yielding four groups: Young_Fast_, Young_Slow_, Old_Fast_, and Old_Slow_. Eligible participants needed to walk without assistance for at least 5 min, have no orthopedic or pain conditions, have no neurological conditions, and have no diagnosed mental health conditions (e.g., anxiety, depression, etc.). All participants were naive to the experimental protocol and had never experienced split-belt walking. The study was approved by the University of Pittsburgh Institutional Review Board. All subjects gave informed consent before testing.

### General Split-Belt Paradigm

To increase the generalization of split-belt patterns [4], all participants performed a gradual split-belt paradigm consisting of 3 experimental epochs: baseline, adaptation, and post-adaptation (Fig. 1A). First, subjects performed a baseline epoch where they walked overground and on the treadmill—this baseline period allowed us to characterize participants’ typical gait when walking overground and on the treadmill. For the overground baseline period, subjects walked continuously back and forth on a 9.2-meter walkway for 6 minutes at their self-selected walking speed. Then, participants performed a baseline period on the treadmill during which subjects walked with both belts moving at the same speed (i.e., tied belts) at mean speed, which depended on the experimental group they were assigned to (see “Experimental groups” section below) for 150 strides. A stride is the time between two consecutive foot landings (i.e., heel strikes) of the same leg. After the baseline condition, subjects experienced the split-belt adaptation condition for two blocks on the treadmill. This adaptation period assessed subjects’ ability to adjust their locomotor patterns to the split-belt environment. In the first adaptation block, subjects experienced a gradual split-belts adaptation where both belts started at their mean walking speed and then linearly diverged for 600 strides. The belt under the participants’ dominant leg started to speed up while the belt on their non-dominant leg started to slow down until the dominant leg was moving twice as fast (2:1 ratio) as the non-dominant leg (see an example of sample subjects in Fig. 1B). Leg dominance was determine based on the self-reported leg used to kick a ball [17]. A catch condition (10 strides) in which participants walked with tied belts at their mean walking speed was allocated right after the first adaptation block. The behavior during this catch period was used to assess the amount of learning in the treadmill context (trained environment). For the second adaptation block, participants experienced an abrupt split-belt perturbation where they walked with their dominant leg moving twice as fast as their non-dominant leg for 300 strides. This block was used to re-adapt participants’ gait to the split-belt environment following the catch. During the second adaptation block, all subjects had a break after the first 150 strides; this was introduced to avoid fatigue and ensure that all participants could complete the experimental protocol. Immediately after the second adaptation block, all participants experienced the overground post-adaptation condition. This condition evaluated the generalization of learned movements to an untrained environment (overground). During the overground post-adaptation, subjects again walked back and forth on a walkway at their self-selected speed for 6 minutes. Subjects were transported from the treadmill to the walkway with a wheelchair to ensure the first steps after adaptation were recorded. Residual context-specific aftereffects were assessed during a treadmill post-adaptation condition, where subjects walked on the treadmill with tied belts for 300 strides at their mean walking speed.

**Figure 1:**
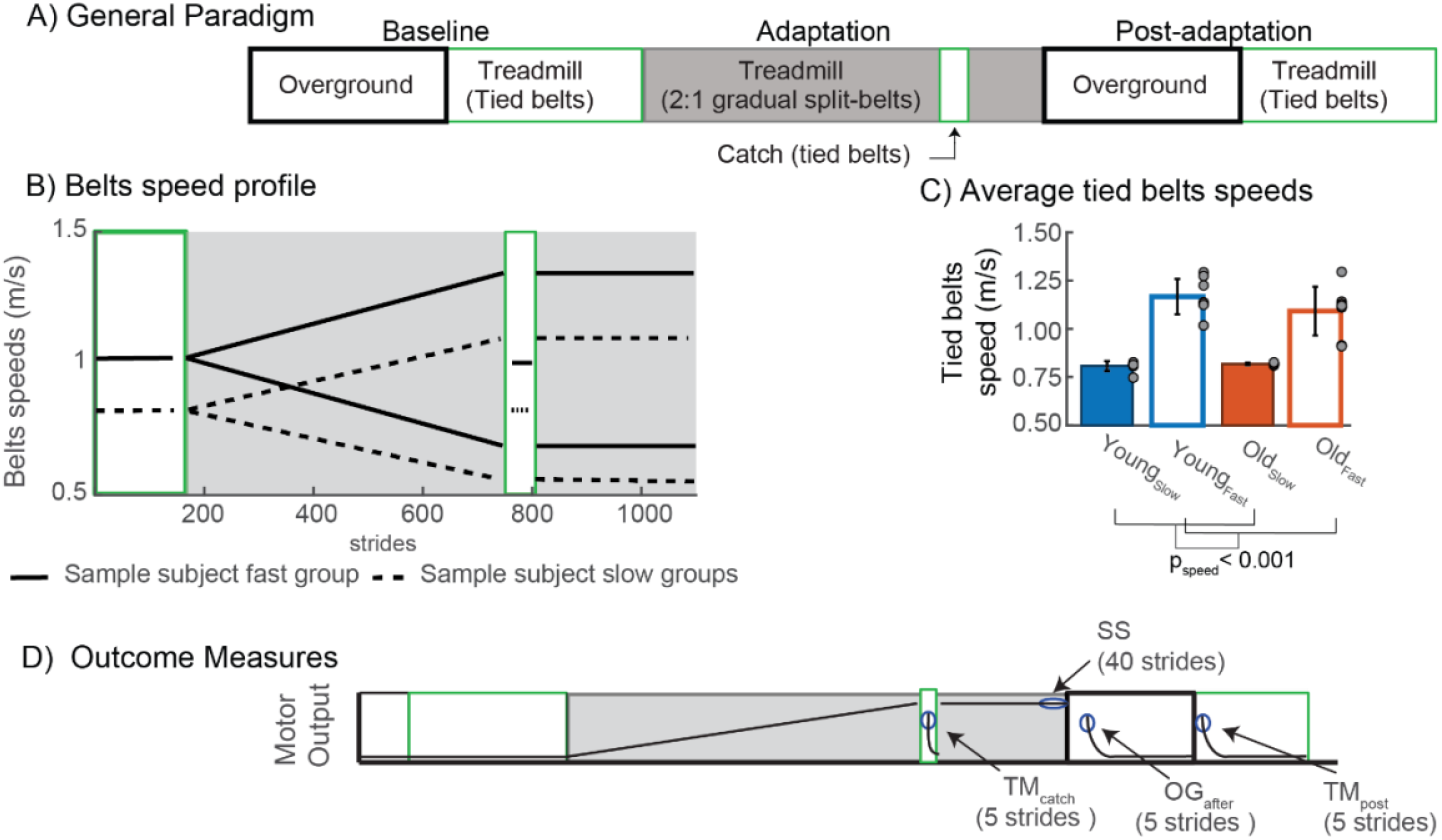
A) Split-belt treadmill paradigm experienced by young and older groups. B) Speed profiles experienced by two example subjects. We illustrate the time course of the speeds experienced by the participants. The speeds on the green rectangle represent the speed during the tied belts (baseline and catch); we can see that the tied belt speeds of these participants were different—Group-averages speed experienced during tied belts. Bar plots represent the group’s average walking speed ± standard error; gray dots represent the individual subjects’ speed during the tied belts condition. The walking speeds for the fast groups is significantly greater than those in the slow groups, as seen by the significant speed effect. D Outcome measures: TM_catch_, SS (Steady State), OG_post,_ and TM_post_ were measured at the periods of interest.

For safety purposes, subjects could hold to a handrail in front of them during the first few steps of the baseline, adaptation, and post-adaptation conditions on the treadmill. They also wore a safety harness while walking on the treadmill; this harness did not support participants’ body weight. A custom-built divider was placed between participants’ legs to prevent them from stepping on the same belt with both legs while walking on the treadmill.

### Experimental groups

Young adults and older adults were randomly assigned to a split-belt paradigm at either a slow or a fast mean walking speed to investigate the effect of walking speed. This mean speed was defined as the average speed between the belts that participants experienced on the split-belt condition on the treadmill during the adaptation epoch. Participants in the slow groups walked at a fixed speed of around 0.75 m/s (Young_Slow_, 5 females, n=8, age: 22.17±1.03 yrs. and Old_Slow_,4 females, n=8, age: 75±6.7 yrs.). We used participants’ comfortable fast walking speed to ensure everybody could complete the task. To measure this, we asked participants to walk on the treadmill and adjust their speed until they found a comfortable fast walking speed. To speed up the treadmill, participants walked farther forward, and to slow down the treadmill participants needed to walk farther back on the treadmill; once they found their fast, comfortable walking speed, they were asked to walk on the middle of the treadmill. Young adults in the fast group (Young_Fast_, 2 females, n=8, age 20±8.67 yrs.) had a mean walking speed of 1.03±0.22 m/s, while the older adults adapted at the fast-walking speed (Old_Fast_, 5 females, n=8, age: 71.2± 3.2 yrs.) had a mean speed 0.99±0.28. The mean walking speeds were not fixed within the groups; however, participants in the fast-walking group walked significantly faster than those in the slow groups (Fig. 1C, F(1, 28) = 128.66, p_speed_ < 0.001), and the mean speeds were not different across age groups (F(1, 28) = 1.33, p_age_= 0.258). Of note, two of the subjects in the old group walked at the 1:1.8 ratio because their fast-walking speed approached the walk-to-run transition. We included their data in the analysis because they did not show differences between their behavior and the rest of the participants in that group. Additionally, data for the catch condition for one subject in the young fast group could not be collected due to technical difficulties. However, the data was included for the rest of the analysis because the behavior of the catch condition is not the primary outcome measurement for this study.

#### Data Collection

Kinematic data were collected to characterize subjects’ behavior. This data was collected at 100 Hz with a passive motion analysis system (Vicon Motion Systems, Oxford, U.K.). Subjects’ movements were assessed through passive reflective markers placed bilaterally on bony landmarks at the ankle (i.e., lateral malleolus) and the hip (i.e., greater trochanter); additional markers were placed asymmetrically on shanks and thighs to differentiate between the legs. Post-processing event detection like heel strike (i.e., foot landing) and toe-off (i.e., foot lifting) was identified with kinematic data [3], [4], [12], [18]. To ensure that participants take the same number of strides on the treadmill, we used a threshold of 30 Newtons to track participants’ heel strikes and toe-off in real-time. Kinetic data were recorded at 1000 Hz with an instrumented split-belt treadmill (Bertec, Columbus, OH).

#### Data Analysis

##### Gait parameters

Subjects’ kinematics were characterized by step length asymmetry (SL_asym_). SL_asym_ is conventionally used to describe gait adaptation and generalization in split-belt studies (e.g., Reisman et al., 2005; Torres-Oviedo and Bastian, 2010; Sombric et al., 2017) as it is of clinical importance (e.g., Jørgensen et al., 1995; Patterson et al., 2008). SL_asym_ was defined as the difference between two consecutive step lengths and normalized by the sum of step lengths (Eq. 1), where a step length (S.L.) is defined as the distance between heel markers at heel strike. SL_Fast_ represents the step length when the fast leg is leading, and SL_Slow_ represents the step length when the slow leg is leading. Thus, a zero value for SL_asym_ indicated that both steps were the same length. By convention, SL_asym_ is positive when the step length of the fast (dominant) leg is longer than the slow (non-dominant) leg. Subjects’ SL_asym_ was normalized by the sum of subjects’ step lengths to account for intrasubject differences in step lengths making all our gait parameters unitless.

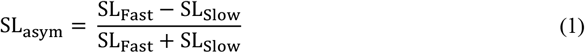

The adaptation of SL_asym_ is due to changes in subjects’ forward and trailing limb position [22], [23]. Previous work has suggested that subjects’ limb position is controlled by different mechanics [23]. Furthermore, changes in inter-limb position asymmetries are associated with changes in subjects’ temporal and spatial walking patterns [18]. Thus, subjects’ asymmetry in the leading leg position (Lead_asym_, Eq. 2) and trailing leg position (Trail_asym_, Eq. 3) were quantified, where Lead_Fast_ and Lead_Slow_ represented the forward position of the leg at fast and slow heel strike, respectively. A positive Lead_asym_ value indicates that the fast leg landed farther from the body than the slow leg. For trialing leg asymmetries, Trail_Fast_ represented the position of the fast leg at the slow strike and vice versa for the Trail_Slow_. By convention, a negative Trail_asym_ value indicates that the fast leg was farther back from our body than the slow leg. Subjects’ limb positions were computed based on subjects’ body position, defined as the average position of the two hip markers.

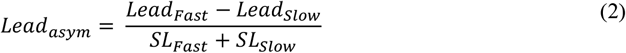

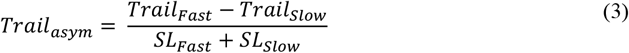

##### Outcome Measures

We analyzed different epochs during the gradual split-belts paradigm to characterize subjects’ adaptation, treadmill aftereffects, and overground aftereffects for each of the parameters defined above (SL_asym_, Lead_asym_, and Trail_asym_) (Fig. 1D). For our analysis, we removed the subjects’ baseline biases for all our outcome measures. This was done by subtracting the baseline-specific (overground or treadmill) values for each gait parameter averaged over the last 40 strides during baseline. In addition, data during the last five strides of each condition were excluded to eliminate effects linked to the treadmillstopping.

We quantified subjects’ steady state (S.S.) to assess adaptation performance. S.S. measures how much participants have changed their walking patterns by the end of the second adaption block, right before they walk overground. We defined S.S. as the mean of the last 40 strides of adaptation.

We computed the average of the initial five strides during three conditions to measure subjects’ aftereffects. First, we calculated the aftereffects during the catch trial (TM_catch_) to assess subjects’ learning during the gradual adaptation. Second, to measure the generalization of adapted movements from the treadmill to the overground walking, subjects’ aftereffects (OG_after_) were assessed during the overground post-adaptation condition. Lastly, to quantify the aftereffects specific to the treadmill environment, the aftereffects on the treadmill during the post-adaptation were assessed (TM_post_). For TM_post,_ large values indicated more contextualization of the split-belt walking patterns, while small values indicated less contextualization. TM_post_ quantifies the amount of contextualization because these aftereffects are measured in the same context as the training environment after overground walking.

#### Statistical analysis

##### Power Analysis

The number of subjects per group was estimated using the overground aftereffects of step length asymmetry from previously published data [12]. We used a between-age-groups variance of 0.00015 and an estimated error variance of 0.0057. An effect size of d=0.53 was obtained from a two-way ANOVA (analysis of variance) of the published data. Our power analysis indicated that an estimated sample size of n=8 subjects per group would allow us to detect this effect size with a significance level of α =0.05 and statistical power of 80%. This number of participants is consistent with existing literature assessing the generalization of locomotor patterns (Torres-Oviedo 2012, Torres-Oviedo and Bastian 2010). This power analysis was done using STATA (StataCorp LLC, TX).

##### Planned analysis

To understand the effect of age and walking speed on the adaptation and generalization of the gait patterns of interest (SL_asym_, Lead_asym_, and Trail_asym_), two-way ANOVAs (analysis of variance) were used. For this, subjects’ age and walking speed were used as independent variables, and our outcome measures (i.e., SS, TM_catch,_ OG_after_, TM_post_) as dependent variables. A post-hoc analysis using Tukey’s HSD correction was performed if a significant interaction was found. The effect size for the significant factor is reported. A significance level α = 0.05 was used for all analyses. STATA was used to perform all statistical analyses (StataCorp L.P., College Station, TX).

## RESULTS

### Young and older adults have similar gait adaptations, but young adults show more treadmill aftereffects than older people

Subjects used similar strategies to update their kinematics to the gradual split-belt. This is shown by an overlap in the groups time courses during the adaptation epoch (Fig. 2A). Furthermore, when we quantified the differences on their steady state at the end of the second adaptation block, we did not observe any significant interactions (SLasym: F(1,28) = 0.17, p = 0.68; Leadasym: F(1,28) = 0.07, p = 0.79; Trailasym: F(1,28) = 0.71, p = 0.41) nor any age effects (SLasym: F(1,28) = 0.00, p = 0.96; Leadasym: F(1,28) = 0.43, p = 0.52; Trailasym: F(1,28) = 0.38, p = 0.54), and there was no effect of walking speed on steady-state values either (SLasym: F(1,28) = 1.13, p = 0.29; Leadasym: F(1,28) = 0.56, p = 0.46; Trailasym: F(1,28) = 0.97, p = 0.33), as illustrated in Fig 2B. Thus, the adaptation of a gradual perturbation is not modulated by subjects’ age or walking speed.

**Figure 2:**
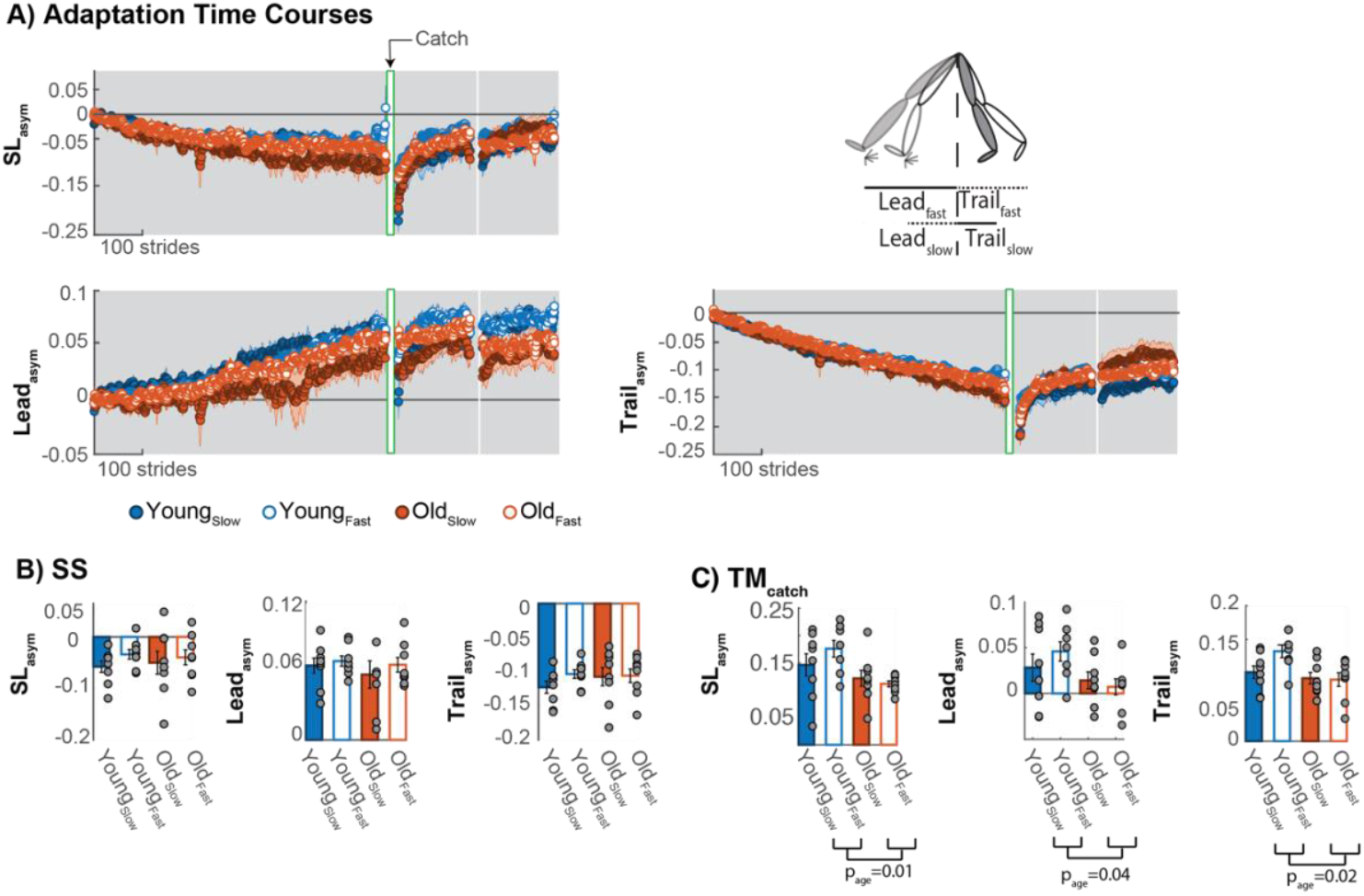
A) Adaptation time courses for groups SL_asym_, Lead_asym,_ and Trail_asym_ during subjects’ adaptation; each dot represents the average of 5 consecutive strides, and the shaded colored areas indicate the standard error for each group. Shaded gray areas represent the adaptation blocks; the green rectangle area represents the catch trial (tied belts), and the white region in the resting break. B) Bar plots indicate the average steady-state behavior per group ± standard error. Gray dots indicate individual subjects. C) Bar plots indicate the average catch behavior per group ± standard error. Gray dots indicate individual subjects’ data.

While all age groups reached a similar adapted state at the end of the adaptation, healthy aging affected their treadmill aftereffects. Young adults’ treadmill aftereffects were larger compared to those of older adults (TM_catch_, Fig. 2C). This is consistent with a significant age effect on subjects’ TM_catch_ aftereffects (SL_asym_, F(1,27)=7.48, p_age_ =0.01). When looking at subjects’ limb position asymmetry, we found that older participants had smaller aftereffects than young groups in both aspects of step length asymmetry: leading leg asymmetry (Lead_asym_, F(1,27)= 4.70, p_age_ = 0.04) and trailing leg asymmetry (Trail_asym_, F(1,27)=6.14, p_age_ = 0.02). However, walking speed did not influence their treadmill aftereffects (SL_asym_, F(1,27)=0.36, p_speed_ =0.56; Lead_asym,_ F(1,27)= 0.20, p_speed_=0.66; Trail_asym_ F(1,27)= 2.03, p_speed_ = 0.17) nor showed a significant interaction (SL_asym_, F(1,27)=1.49, p_age#speed_ =0.23; Lead_asym,_ F(1,27)= 1.03, p_age#speed_=0.32; Trail_asym_ F(1,27)= 2.63, p_age#speed_ = 0.12). Thus, our results suggest that healthy aging reduces the human capacity to learn a new walking pattern.

### Walking speed regulates the generalization of walking patterns from split-belt treadmill to overground in an age-specific manner

Subjects’ generalization of split-belt walking patterns depends on their age group and walking speed during training. Fig. 3A shows the time courses of subjects’ overground aftereffects. Notice that younger adults generalize more the split-belt walking when they walk faster, while old adults generalize more of these walking patterns when walking slower. A significant interaction in subjects’ SL_asym_ between subjects’ age and walking speed further supports these findings (F(1,28)= 19.25, p_age#speed_ < 0.001, Fig. 3B). A post hoc analysis showed that young adults adapted at a fast-walking speed carryover more than those in the slow walking speed group (Young_Fast_ vs. Young_Slow_, p=0.02) and older adults adapted at fast walking speed (Old_Fast_ vs. Young_Fast_, p=0.02). Additionally, older adults adapted at a slower walking speed carryover more than those adapted at a faster speed (Old_Slow_ vs. Old_Fast_, p=0.02), and young adults adapted at a slow speed (Yong_Slow_ vs. Old_Slow_, p=0.02). Importantly, we did not find significant differences between young adults whoadapted at a fast-walking speed and old adults who adapted at aslow walking speed (Yong_Slow_ vs. Old_Slow_, p=1.0). Thus, healthy aging and treadmill walking speed modulate the generalization of subjects’ step length asymmetry.

**Figure 3:**
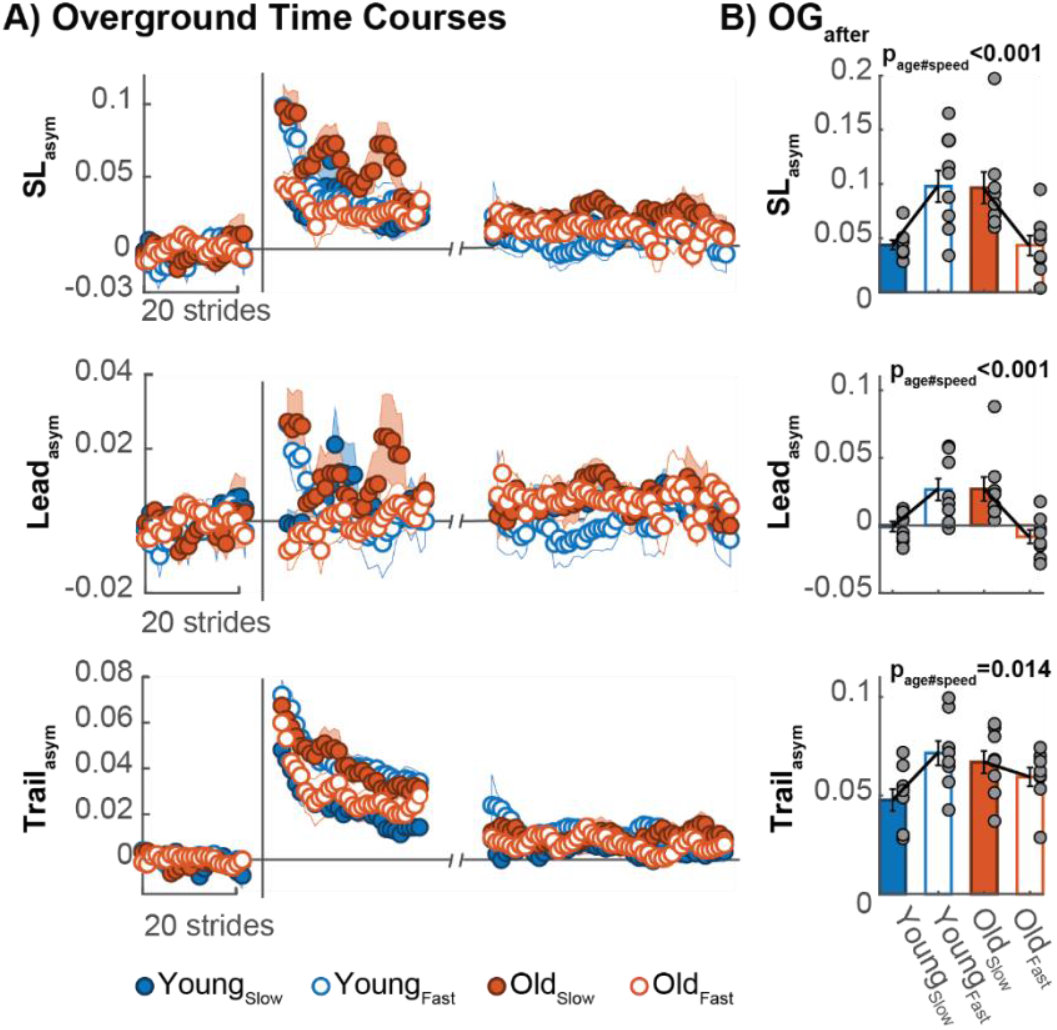
A) Group time course of step length asymmetry during the overground baseline and post-adaptation. Dots represent the average of 5 consecutive strides, and the shaded colored regions indicate group standard errors. B) Bar plots indicate OG_after_ (averaged values for each group) ± standard errors. Gray dots indicate individual subjects’ behavior. The black lines represent an interaction effect. For example, notice that young adults generalize more when they walk at a faster speed while older adults generalize more at a slower speed.

When we analyzed subjects’ limb position, we found a significant interaction for the leading leg asymmetry (F(1,28) =19.06, p_age#speed_ < 0.001) and trailing leg asymmetry (F(1,28)= 6.95, p_age#speed_ = 0.014). For the leading limb position, a post hoc analysis showed that old adults walking at a slow speed generalize more than young adults walking at a similar speed (Old_Slow_ vs. Young_Slow_, p=0.049). Additionally, older adults (Old_Slow_ vs. Old_Fast_, p=0.008) and young adults (Young_Fast_ vs. Old_Fast_, p=0.01) walking at a fast speed placed their limbs farther forward compared to the older adults walking faster. Likewise, a post hoc analysis of subjects’ trailing limb placement showed those in the fast group placed their slow leg farther behind their body than those in the slow group (Young_Fast_ vs. Young_Slow_, p=0.039). Thus, subjects’ age and walking speed modulate the generalization of the leading and trailing legs’ position.

### Walking speed modulates the washout of context-specific walking patterns

We found that walking speed has an effect on the resilience of context-specific motor memories developed on the split-belt treadmill. Figure 4 shows subjects’ remaining aftereffects when returning to the treadmill after overground walking. When quantifying the differences in subjects remaining treadmill aftereffects, we found that healthy aging does not affect the subjects remaining aftereffects (SL_asym_:F(1,28)=0.01, p_age_ = 0.90; Lead_asym_: F(1,28)=0.11, p_age_= 0.75;Trail_asym_: F(1,28) = 0.30, p_age_=0.59). However, subjects walking speed does affect the remaining aftereffects SL_asym_ (F(1,28)= 6.58, p_speed_ = 0.02), where those adapted at a slower speed have larger remaining aftereffects when returned to the treadmill. Noticed that, when looking at subjects leading and trailing leg asymmetry, we found that neither parameter directly drives this difference (Lead_asym_: F(1,28)= 1.53, p_speed_=0.23; Trail_asym_: F(1,28)= 2.40, p_speed_=0.13) nor showed a significant interaction (SL_asym_:F(1,28)=0.75, p_age#speed_ =0.39; Lead_asym_: F(1,28)<0.01, p_age#speed_= 0.98;Trail_asym_: F(1,28) = 2.07, p_age#speed_=0.16). Thus, our results indicate that slower walking during adaptation and post-adaptation leads to larger aftereffects on the treadmill (training context).

**Figure 4:**
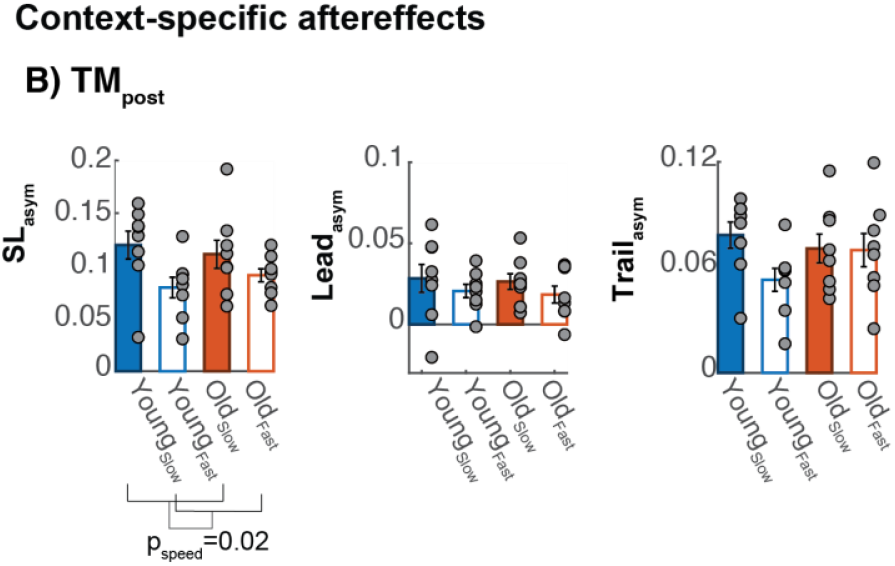
Treadmill post-adaptation: Bar plots indicate the group average remaining aftereffects on the treadmill (training context) following the overground walking ± standard errors. Dots showed individual subjects’ behavior. Horizontal lines represent a significant difference between the group speeds.

## DISCUSSION

### Summary

We investigated how walking speed and healthy aging affect participants’ ability to adapt and generalize their walking patterns. As anticipated, we found an interaction between healthy aging and walking speed in subjects’ generalization of movements. In other words, we observed that young adults generalize more whenadapted at a fast-walking speed, while older adultsgeneralize more whenadapted at a slow walking speed. Interestingly, this interaction was not observed in the training context (treadmill), where we found either an age effect (during the catch) or a speed effect on subjects’ aftereffects (following overground). In sum, we found that healthy aging and walking speed modulate the generalization and formation of context-specific walking patterns.

### Walking speed regulates age-specific transfer of split-belt walking patterns and treadmill-specific motor memories

We found an interaction between subjects’ age and walking speed on the transfer of walking patterns from the treadmill to overground. In other words, older adults transferred more their walking patterns across walking conditions when adapted at slower speeds, whereas young adults did so more when adapted at faster speeds (Fig. 3). Previous studies have shown that decreased contextual differences between task conditions improve the transfer of movements across walking environments [3], [11], [24]–[27], which supports the contextual inference theory [28] positing that larger deviations from participants’ regular movements (e.g., differences between training and preferred walking speeds) facilitate the context-specificity of motor memories, thereby, reducing their generalization. Thus, we can interpret that walking on the treadmill at a speed closer to one’s preferred speed reduces the contextual differences between the training (treadmill) and testing (overground) environments. This interpretation is supported by previous work showing that young adults’ self-selected speed is faster than older adults’ self-selected speed [13]–[15], [29] and faster self-selected pace during overground walking leads to a faster-preferred speed on the treadmill [30]. Thus, it is reasonable to speculate that the slow speed during treadmill adaptation is closer to the self-selected speed of older adults, whereas the fast speed during adaptation is more comfortable for younger adults. Future studies are needed to determine if the generalization of walking patterns follows an inverted U-shape generalization function [31]–[33], where the maximum amount of generalization corresponds with subjects’ self-selected speed and the farthest away from that speed, the less carry over between the training and testing environments.

Walking speed also regulated the washout of treadmill-specific aftereffects post-adaptation. However, this speed effect did not reflect the interaction that we found between age and walking speeds during overground walking. In other words, the groups that transferred more of their split-belt motor pattern did not wash out more of the treadmill-specific motor memory. This lack of correspondence between the transfer and washout of device-induced motor memories has been reported in previous generalization studies [4], [12], [16], [18], which suggests that there are distinct factors mediating the transfer and washout of device-induced motor memories. Specifically, we found that young and older adults walking at slower speeds exhibited larger remaining aftereffects when returning to the treadmill after overground walking (Fig. 4).

These results can be interpreted in two ways. One interpretation is that slow walking elicits larger treadmill aftereffects than fast walking [34]. Therefore, the magnitude of aftereffects on the treadmill is a function of the speed at which these aftereffects are tested, regardless of the speed at which they are trained. Another interpretation is that overground walking affected the amount of remaining aftereffects. Previous study shows that slow walking washes out motor memories tested at fast and slow walking speeds, whereas fast walking only washes out motor memories tested at fast, but not slow, walking speeds [34]. Consistently, overground walking occurring at a wide speed range could have washout motor memories formed at fast walking speeds but not those formed at slow walking speeds. Lastly, we cannot disregard the participant’s walking speed during training; thus, slow walking in a novel environment may lead to more resilient motor memories than fast walking. However, future studies will be needed to parse out the effect of the training speed and testing speed on the participants’ treadmill-specific aftereffects.

### Healthy aging reduces the sensorimotor aftereffects to a gradual perturbation but not the ability to counteract the novel split-belt environment

We showed that younger adults have larger aftereffects than older adults when assessed on the treadmill during the catch trial (Fig. 2C). This observation is at odds with previous work showing that sensorimotor recalibration, as quantified by treadmill aftereffects, is not affected by aging (walking: Bruijn et al., 2012; Malone and Bastian, 2016; Sombric et al., 2017; Iturralde and Torres-Oviedo, 2019; Sombric and Torres-Oviedo, 2021) and reaching: Buch et al., 2003; Bock, 2005; Bock and Girgenrath, 2006; Seidler, 2007). The development of locomotor memories is mediated by two factors, exposure to larger errors during the initial adaptation and the duration of the full perturbation [42]. Notice that we measured the subjects’ treadmill aftereffect after the ramp phase (Fig. 1D), meaning that no larger errors were providing salient cues about the changes in the environment, and participants did not experience the full split-belt perturbation. Previous work showed that healthy aging increases sensory [43]–[45] and motor noise [46]–[49], which may reduce participants’ sensitivity to errors [12], [50]. This reduction in their sensitivity to errors can impact older adults’ ability to recalibrate movements in a gradual perturbation where only small errors are present. Thus, healthy aging reduces sensorimotor aftereffects induced by gradual perturbations.

On the other hand, neither healthy aging nor walking speed affects the adaptation to the split-belt perturbation. Namely, subjects had similar steady-state behavior at the end of the adaptation (Fig. 2B). This indicates that participants’ ability to counteract the perturbation was not affected by their age or walking speed. While these observations are consistent with previous work [12], [37], it is at odds with other findings suggesting that speed [51] or healthy aging [36], [52]–[55] affect participants’ steady-state behavior. We can reconcile our results with previous findings if we consider the differences in methodologies. For example, participants in our protocol experience large errors several times (i.e., after the catch and after each resting break), whereas previous split-belt studies [36], [51] only experience large errors once upon introduction of the split-belt perturbation. These differences are relevant since large errors lead to more adaptation [56], [57], meaning that large errors after the catch trial and the break help participants counteract the perturbation better. Therefore, older adults adapt their gait similarly to young adults when multiple large errors are experienced during the adaptation, regardless of their treadmill speed.

Furthermore, previous work shows that older adults rely on explicit strategies to perform sensorimotor tasks [16], [58], [59]; however, these explicit strategies do not affect the recalibration of movements [60]. In other words, while older adults need salient cues about the environmental changes to elicit explicit strategies to update their motor patterns, which will allow them to reach a similar steady state as younger adults, these explicit strategies would not translate into more internal recalibration of movements, quantified by treadmill aftereffects. Thus, explicit strategies can aid participants’ ability to counteract a perturbation, but it does not help with the formation of a novel motor memory.

## CLINICAL IMPLICATIONS

The generalization of movements from trained to untrain situations is critical for rehabilitation. It has been shown that split-belt walking can improve the gait of post-stroke survivors [2], [61], and these changes persist with multiple exposures [62], [63]. However, not everything learned in the training context carries over to day-to-day activities[1], [62]. Thus, understanding how to make the movement more generalizable across different situations is relevant for rehabilitation. Our results suggest that training participants close to their preferred walking speeds during rehabilitation therapies might lead to more general movements, whereas training at uncomfortable walking speeds may tie the learned movement to the training context. Additionally, we found that older adults do not exhibit treadmill-specific aftereffects as quickly as young adults in response to gradual perturbations. However, large device-induced perturbations enable older adults to reach similar sensorimotor adaptation as younger adults without affecting the transfer of sensorimotor patterns. Thus, it may be beneficial for older patients to experience larger errors while developing new motor memories since large errors during locomotor adaptation can improve the generalization of their movements[5].

## ETHICS STATEMENT

This study was carried out following the recommendations from the University of Pittsburgh Institutional Review Board with written informed consent from all subjects. In addition, all participants gave written informed consent following the Declaration of Helsinki.

## AUTHOR CONTRIBUTIONS

D.M. contributions include processing and analysis of the data, writing a complete draft of the manuscript, revising work for important intellectual content, final approval of the version to be published, and agreement to be accountable for all aspects of the work. C.S. contributions include data acquisition and processing, interpretation of the data and final approval of the version to be published, and agreement to be accountable for all aspects of the work. Finally, G.TO contributions include the conception and design of the work, revising the work, final approval of the version to be published, and agreement to be accountable for all aspects of the work.

